# A cross-species neural integration of gravity for motor optimisation

**DOI:** 10.1101/728857

**Authors:** Jeremie Gaveau, Sidney Grospretre, Dora Angelaki, Charalambos Papaxanthis

## Abstract

Recent kinematic results, combined with model simulations, have provided support for the hypothesis that the human brain uses an internal model of gravity to shape motor patterns that minimise muscle effort. Because many different muscular activation patterns can give rise to the same trajectory, here we analyse muscular activation patterns during single-degree-of-freedom arm movements in various directions, which allow to specifically investigating gravity-related movement properties. Using a well-known decomposition method of tonic and phasic electromyographic activities, we demonstrate that phasic EMGs present systematic negative phases. This negativity demonstrates that gravity effects are harvested to save muscle effort and reveals that the brain implements an optimal motor plan using gravity to accelerate downward and decelerate upward movements. Furthermore, for the first time, we compare experimental findings in humans to monkeys, thereby generalising the Effort-optimization strategy across species.

## Introduction

The ability to purposely move one’s own body is a critical survival function that humans and animals master with apparent ease. However, even the most straightforward body limb movement entails inherent difficulties for which the motor system has evolved sophisticated solutions (Franklin and Wolpert, 2011; Shadmehr and Wise, 2005; Shadmehr et al., 2010; Wolpert and Ghahramani, 2000). Amongst others, one of these solutions is to learn and store internal models to disambiguate sensory information and to predict forthcoming movement dynamics. On earth, a pervasive component affecting perception and motion is gravity (Bringoux et al., 2012; Dakin and Rosenberg, 2018; Elmore et al., 2014; Gaveau et al., 2011, 2016; Jorges and Lopez-Moliner, 2017; Klein et al., 2018; Lacquaniti et al., 2013; McIntyre et al., 2001; Van Pelt et al., 2005; Rosenberg and Angelaki, 2014; Saradjian et al., 2014; La Scaleia et al., 2019; Senot et al., 2005; Tajadura-Jimenez et al., 2018; De Vrijer et al., 2008). Studies in both humans and non-human primates have provided strong evidence that the brain evolved an internal representation of gravity. This representation is thought to involve neural computations of the brain stem, the cerebellum, the vestibular cortex, and the anterior thalamus (Angelaki et al., 2004; Indovina et al., 2005; Laurens et al., 2013a, 2016; Miller et al., 2008; Rousseau et al., 2016). Although the neural representation of gravity is well documented, how it may benefit the production of suitable motor commands is unclear. Yet, living organisms produce successful movements while facing gravity effects every day. It is of critical importance to shed light on the neural computations that underpin motor planning and control in the gravity field.

When moving our body limbs, the brain must generate neural commands that consider both inertial forces and gravity forces. Functional segregation of the inertial forces – related to the velocity and acceleration of the limb – and the gravity forces – related to the position of the limb – was long assumed (Atkeson and Hollerbach, 1985; Flanders and Herrmann, 1992; Hollerbach and Flash, 1982). According to this assumption, the internal model of gravity is used to compensate for the gravity force throughout the entire movement. That is, neural commands produce a muscular force that is equal and opposite to the gravity force. Such a neural policy is thought to facilitate the production of accurate movements to changing directions, amplitudes, durations, and loads, by merely scaling the inertial-dependent part of the motor command. Albeit based upon old literature, this influential *Compensation hypothesis* still guides current research in various fields such as motor control (d’Avella et al., 2008; Guigon et al., 2007; Kadmon Harpaz et al., 2014; Olesh et al., 2017; Russo et al., 2014), movement perception (Cook et al., 2013; Edey et al., 2019) or neuro-rehabilitation (Krabben et al., 2012; Prange et al., 2009a, 2009b, 2012; Raj et al., 2019).

On the other hand, recent kinematic results challenged this prevalent theory by revealing velocity profiles for mono-articular arm movements whose temporal structure changes according to movement direction relative to the gravity vertical (Gaveau and Papaxanthis, 2011; Gaveau et al., 2014, 2016; Gentili et al., 2007; Le Seac’h and McIntyre, 2007). This finding is straightforward because, during mono-articular movements, only the gravity force changes with movement direction; inertial forces are direction-independent. Thus, this observation contradicts the fundamental premise of the *Compensation hypothesis*, which assumes direction-invariant kinematics. In contrast, the reported direction-dependent kinematics of human single-joint rotations are consistent with an optimal control strategy that discounts muscle effort (Berret et al., 2008; Crevecoeur et al., 2009; Gaveau et al., 2014, 2016). Such *Effort-optimization hypothesis* assumes an internal model of gravity to take advantage of its effects rather than to compensate for them.

During adaptation to microgravity, the properties of arm movements further supported this *Effort-optimization hypothesis*. As predicted by an optimisation model, the direction-dependence of arm movements progressively vanished in microgravity (Gaveau et al., 2016). This contrasts to the traditional compensation view that the brain uses internal models of perturbing forces for their compensation, such that stereotypic trajectories can be maintained (Atkeson and Hollerbach, 1985; Hollerbach and Flash, 1982; Shadmehr and Mussa-Ivaldi, 1994). A more recent view is that motor adaptation constitutes a re-optimisation process whereby newly constructed/calibrated internal models generate newly shaped trajectories (Gaveau et al., 2016; Izawa et al., 2008; Selinger et al., 2015; Snaterse et al., 2011).

Support for the E*ffort-optimization hypothesis* is so far limited to kinematic findings. This limitation is problematic because of the redundancy between the muscular level and the kinematic level (Bernstein, 1967). Because many different muscular activation patterns can give rise to the same trajectory, using kinematic data exclusively to infer central processes is insufficient (Burdet et al., 2001; Hagen and Valero-Cuevas, 2017). Establishing that direction-dependent kinematics truly reflects motor commands that discount muscle effort requires additional evidence from muscle dynamics. Here we analyse muscular activation patterns during single-degree-of-freedom arm movements performed in various directions. Furthermore, for the first time, we compared experimental findings in humans to monkeys, thereby generalising the *Effort-optimization* strategy across species.

## Results

We trained three rhesus monkeys to perform earth-vertical and earth-horizontal arm movements around the shoulder joint (Figure 1A). We also asked two groups of humans to perform earth-vertical and earth-horizontal arm movements from two different body orientations (seated upright and 90° tilted in roll). Arm movements were parallel to the participants’ head/feet body axis in a first group (ego-parallel group, n=8, Figure 1B) and perpendicular to it in a second group (ego-perpendicular group, n=8, Figure 1C). Comparison between the ego-parallel group and the ego-perpendicular group allows dissociation of arm movement direction in body-centred and gravity-centred frames of reference.

**Figure 1.**
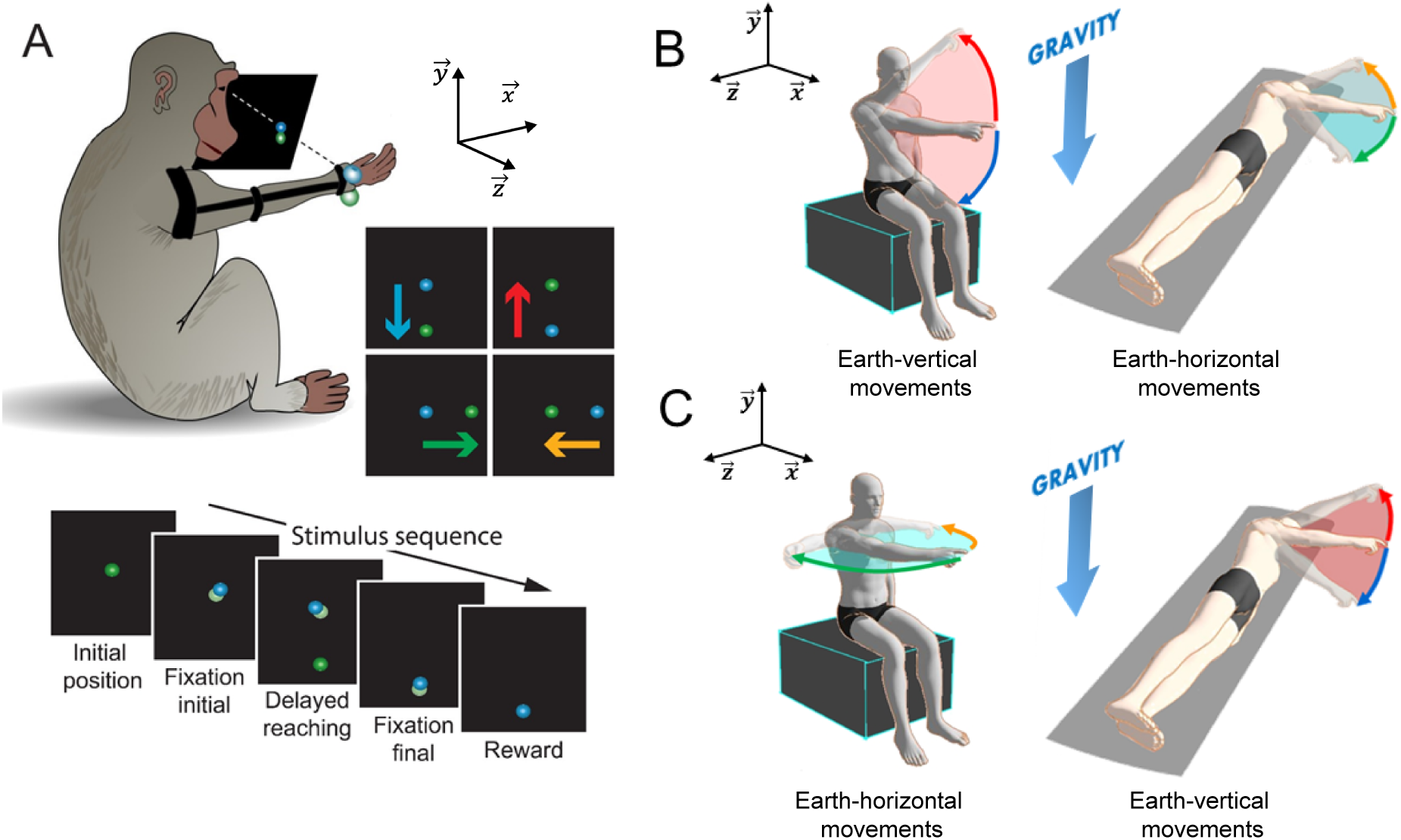
Experimental setup. **A**. Illustration of the monkey task. Using a virtual reality environment, three monkeys performed mono-articular point-to-point arm movements (shoulder rotations, a brace restraining elbow and wrist joints) between sets of two targets. Positions of initial and final targets implied a leftward (yellow arrow), a rightward (green), a downward (blue) or an upward (red) movement. The monkeys performed a delayed-reaching task with fixation periods on the initial and the final target. The lower part of the panel depicts the stimulus sequence. First, the initial target appeared and its colour changed to light green when the monkey’s hand touched it. After a random delay time, the final target appeared, and the monkey had to reach it as soon as the initial target disappeared (additional random delay time). Finally, the monkey obtained a liquid reward after remaining stationary on the final target for another random delay time. **B**. Illustration of the human task for the ego-parallel group. Starting with their right arm perpendicular to the trunk (dark middle arm), height humans performed single-degree-of-freedom reaching arm movements (shoulder rotations) between sets of two targets. Positions of targets implied movements toward the head (shoulder flexion) or the feet (shoulder extension). Participants experienced two conditions of body orientations to record movements in the earth-vertical plane (participant seated, red/blue arrows) and in the earth-horizontal plane (participant reclined, yellow/green arrows); i.e. targets were rotated with the participant. **C.** Illustration of the human task for the ego-perpendicular group (height other human participants, same organisation as the first group, see methods). The positions of targets implied movements toward the right side of the body (shoulder abduction) or the left side of the body (shoulder adduction). Participants experienced two conditions of body orientations to record movements in the earth-vertical plane (participant reclined, red/blue arrows) and in the earth-horizontal plane (participant seated, yellow/green arrows); i.e. targets rotated with the participant. In all three panels, the x, y, z axes illustrate the world coordinate system. For all figures, the colour-code is gravity-centred, as follows: red is against gravity (towards positive Y-axis); blue is with gravity (negative Y), yellow is perpendicular to gravity, leftwards (negative Z); green is perpendicular to gravity, rightwards (positive Z).

### Monkey and human arm movement kinematics follow similar direction-dependent asymmetries

Reaching arm movements typically exhibit bell-shaped velocity profiles (Kelso et al., 1979). Velocity first rises to a peak (acceleration phase) and then declines back to zero (deceleration phase, see Figure 2A-B). As previously reported, single-degree-of-freedom human arm movements show direction-dependent asymmetries in the earth-vertical but not the earth-horizontal plane (Gentili et al., 2007; Le Seac’h and McIntyre, 2007). A shorter and steeper acceleration profile for upward than for downward movements characterises these direction-dependent asymmetries (rise to peak velocity in Figure 2A-B, top traces). Because previous work already demonstrated that this effect was independent of body orientation, Figure 2 presents averaged results across both groups of humans (Fig 2, Suppl. Table 1 and Figure 2, Suppl. Table 2 provides results and statistics for each group). We found that monkeys also exhibit direction-dependent arm kinematics in the vertical plane only (Figure 2A-B, bottom traces). As in humans, the acceleration duration is shorter and steeper for upward than for downward movements.

**Figure 2.**
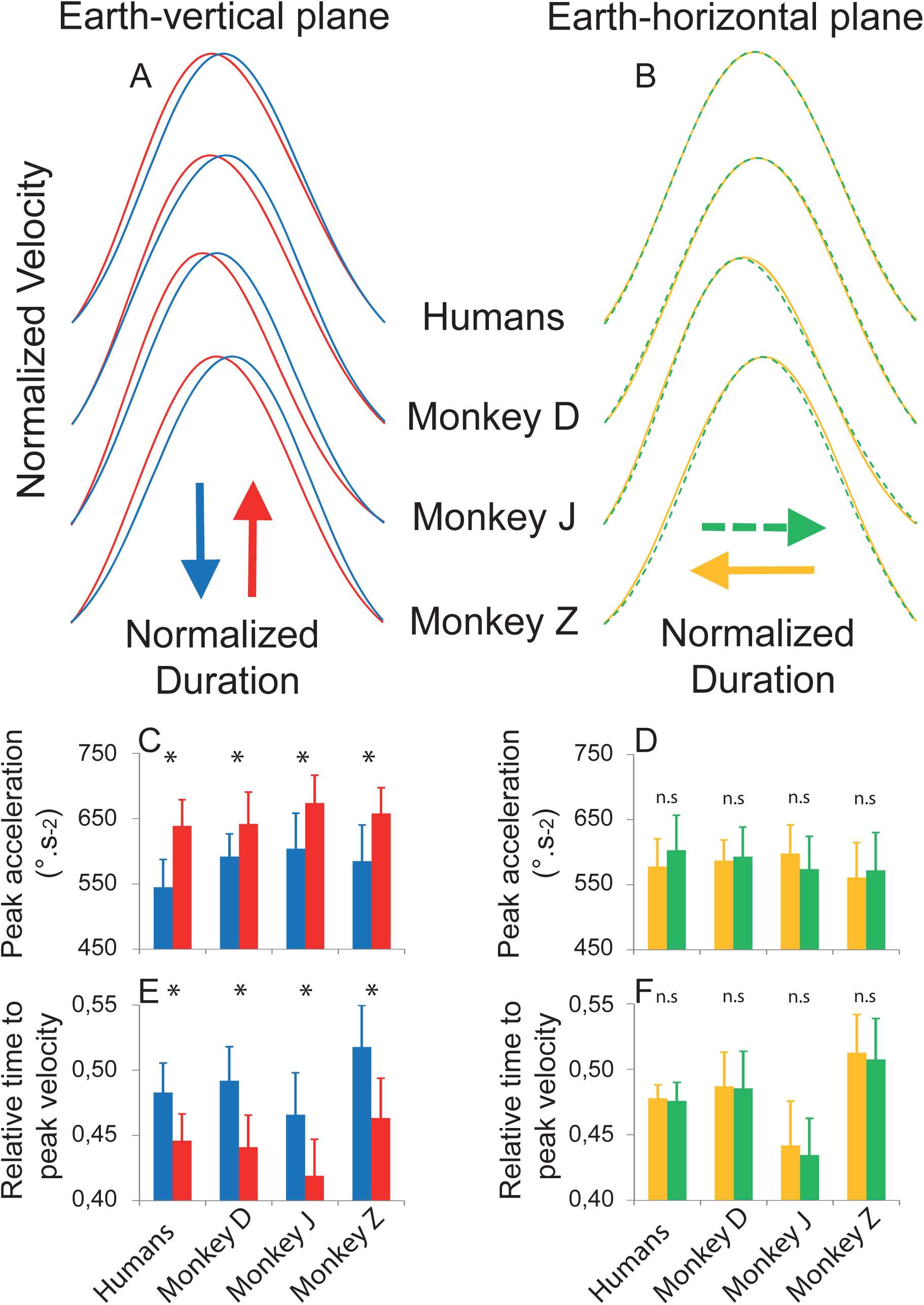
Similar direction-dependent arm kinematics in monkeys and humans. Mean velocity profiles recorded in opposed directions in the earth-vertical (**A**) and the earth-horizontal plane (**B**) for each monkey and all humans (n=16). Coloured arrows indicate movement directions. Traces are amplitude and duration normalised to ease directional comparisons. Direction-dependent kinematics is observed in the earth-vertical plane, but not in the earth-horizontal plane. The amplitude of the acceleration peak for each monkey and all humans (n=16) is presented for opposed directions in the earth-vertical plane (**C**) and the earth-horizontal plane (**D**). The relative duration to peak velocity for each monkey and all humans (n=16) is presented for opposed directions in the earth-vertical plane (**E**) and the earth-horizontal plane (**F**). Error bars represent the standard deviation of the mean between recording sessions for each monkey and between participants for humans.

This motor behaviour was quantified using multiple parameters (see Fig. 2, Suppl. Figure 1A and Fig. 2, Suppl. Table 1). The peak acceleration (which quantifies the steepness of the acceleration phase) and the duration to peak velocity (which specifies the length of the acceleration phase) illustrate this result in Figure 2C-F. Although movement duration and amplitude were similar for all directions (P>0.19 in all cases; see Fig. 2, Suppl. Table 2 for all statistical comparisons), a significant main effect of direction was observed for peak acceleration, peak velocity, relative time to peak acceleration, and relative time to peak velocity (P<0.003 in all cases). Post-hoc comparisons, between opposite directions, yielded significant effects in the earth-vertical but not the earth-horizontal plane (P<0.02 in all cases). Thus, kinematic asymmetries in monkeys are identical to those previously known in humans.

Theoretical models minimising muscle effort predict these asymmetries (Berret et al., 2008; Crevecoeur et al., 2009; Gaveau et al., 2014, 2016). According to such models, the brain takes advantage of gravity effects to accelerate the arm downwards and decelerate the arm upwards. Next, we examine muscular activation patterns to understand the production of direction-dependent arm kinematics further. Because of the redundancy between the muscular level and the kinematic level (Bernstein, 1967), many different muscular activation patterns can give rise to the same velocity profile (Burdet et al., 2001; Hagen and Valero-Cuevas, 2017). Does the neural integration of gravity truly blossom into muscular activation patterns that discount muscle effort?

### Phasic EMG activity supports the Effort-optimization hypothesis

Two types of muscles can contribute to the motion of the arm in the earth-vertical plane: those that pull against gravity (towards positive y-axis in Figure 1) and those that pull with gravity (towards negative y-axis). Hereafter, the first type is named antigravity muscles, and the second type is named gravity muscles. In the earth-horizontal plane, to produce the arm motion, the muscles pull perpendicularly to gravity. Hereafter, the muscles pulling towards the positive z-axis are named rightward-muscles, and the opposite ones are named leftward-muscles.

We applied a simple and widely-used decomposition method to isolate the tonic (gravity-dependent force) and the phasic (inertial-dependent force) EMG components from the full EMG signal (Buneo et al., 1994; d’Avella et al., 2006, 2008; Flanders and Herrmann, 1992; Flanders et al., 1994, 1996; Olesh et al., 2017; Prange et al., 2009b, 2012; Russo et al., 2014). The tonic component emanates from the motionless rest-periods before and after the movement (Fig. 2 Suppl. Figure 1 B, D). The phasic component results from the subtraction of the tonic activity from the total EMG (Fig. 2 Suppl. Figure 1 C-E). This subtraction stems from the *Compensation hypothesis* according to which the tonic would compensate for the gravity force, whereas the phasic would produce the change in velocity (Atkeson and Hollerbach, 1985; Flanders and Herrmann, 1992; Hollerbach and Flash, 1982). Here we take advantage of this method to test straightforward predictions of both the *Compensation* and the *Effort-optimization* hypotheses.

The straightforward prediction from the *Compensation hypothesis* is that phasic EMG activity of antigravity muscles is always positive when the arm is moving. That is, for antigravity muscles to compensate for the gravity force, the total activity in antigravity muscles should be at least equal to the tonic level over the entire movement amplitude. The straightforward prediction from the *Effort-optimization hypothesis* is that the phasic component of antigravity muscles exhibits periods of negativity. Inevitably, for gravity force to assist the arm motion, the antigravity muscle activity should drop below the tonic level that would be required to compensate it.

In accord with the *Effort-optimization hypothesis*, negative periods were frequent for antigravity muscles (Figure 3A, B; purple). The negativity of antigravity muscles precisely occurred when gravity effects could assist movement; that is, during the acceleration of downward movements (Figure 3A) and the deceleration of upward movements (Figure 3B). During those periods, the arm presumably falls free. Gravity muscles, whose effects can only add up to those of gravity, did not exhibit such negative periods (Figure 3C, D). Nor did any muscles for Earth-horizontal movements during which the task requires to compensate for gravity effects (Figure 4). These qualitative results indicate that the presence of negative phasic EMG is specific to the earth-vertical plane and, even more specific, to the timing when gravity can assist movement. Further, it is independent of body-orientation, and both species share it.

**Figure 3.**
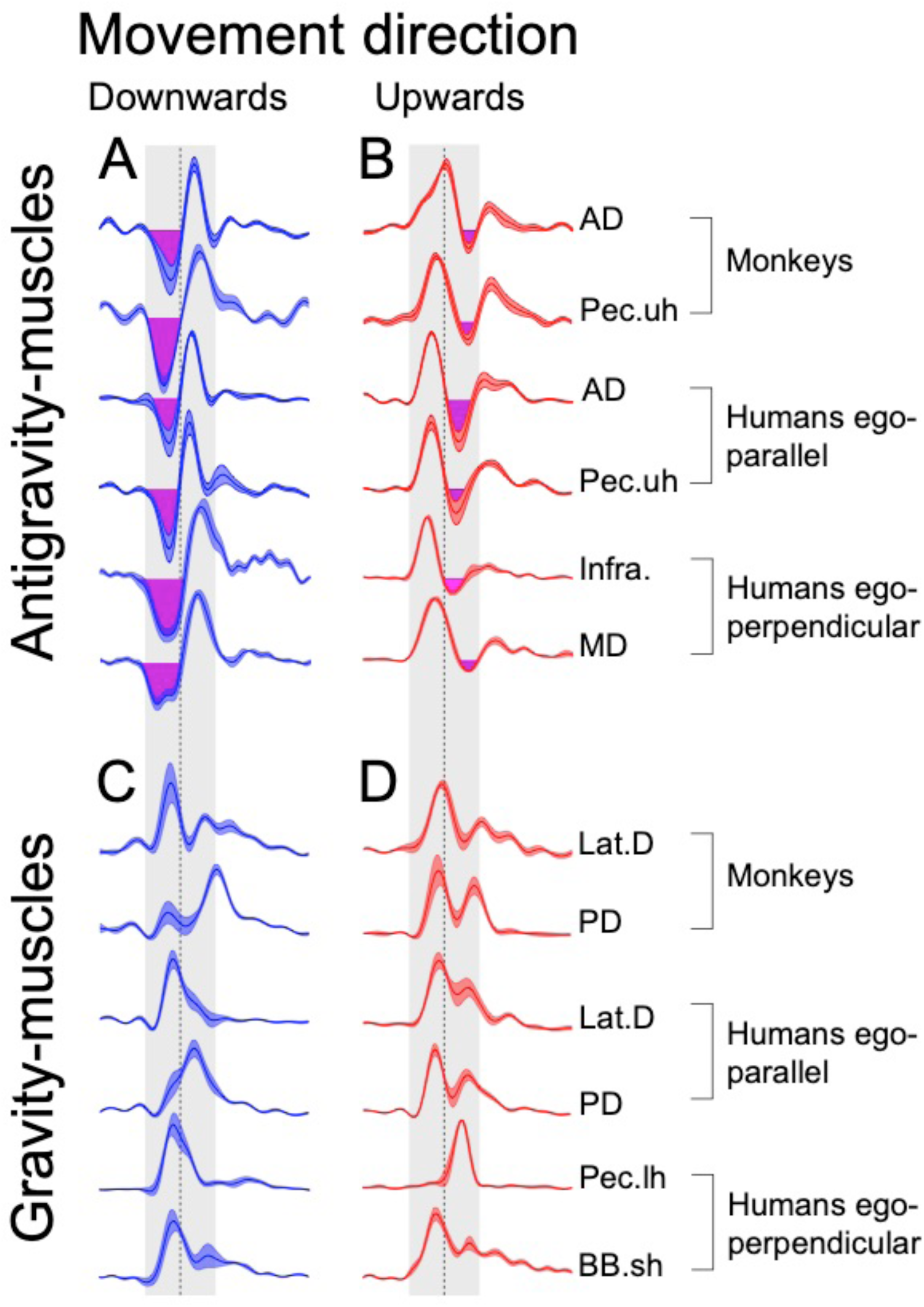
Phasic EMG patterns of earth-vertical movements reveal gravitational force contribution to the motion of the arm. Mean (± SE) phasic EMGs recorded in monkeys (n=3) and both groups of humans (n=8 in each group) during earth-vertical movements. Left column (panels A and C, blue traces) presents EMGs recorded during downward movements while right column (panels B and D, red traces) presents EMGs recorded during upward movements. Upper row (panels A and B) presents EMG activations of antigravity muscles (pulling upwards, against gravity): Anterior Deltoid (AD), upper head of Pectoralis major (Pec. uh), Infraspinatus (Infra.), Middle Deltoid (MD). Those muscles pull against gravity, i.e. away from the final target during a downward movement and towards the final target during an upward movement. Lower row (panels C and D) presents EMG activations of gravity muscles (pulling downwards, with gravity): Latissimus dorsi (Lat.D), Posterior Deltoid (PD), lower head of Pectoralis major (Pec.lh), the short head of Biceps Brachii (BB.sh). Those muscles pull with gravity; i.e. towards the final target during a downward movement and away from the final target during an upward movement. Purple Area denotes phases where epochs of negativity were detected (see Methods). Such epochs were precisely observed for antigravity muscles (Panels A and B) during the acceleration of a downward movement and during the deceleration of an upward movement; i.e., when gravity effects can help the muscle and therefore discount muscle effort. EMG traces were aligned on movement onset and normalised in duration and amplitude before averaging between participants. The grey earth-vertical area denotes movement duration (shifted 100ms backwards to account for the electromechanical delay), and the dashed earth-vertical line denotes 50% of movement duration.

**Figure 4.**
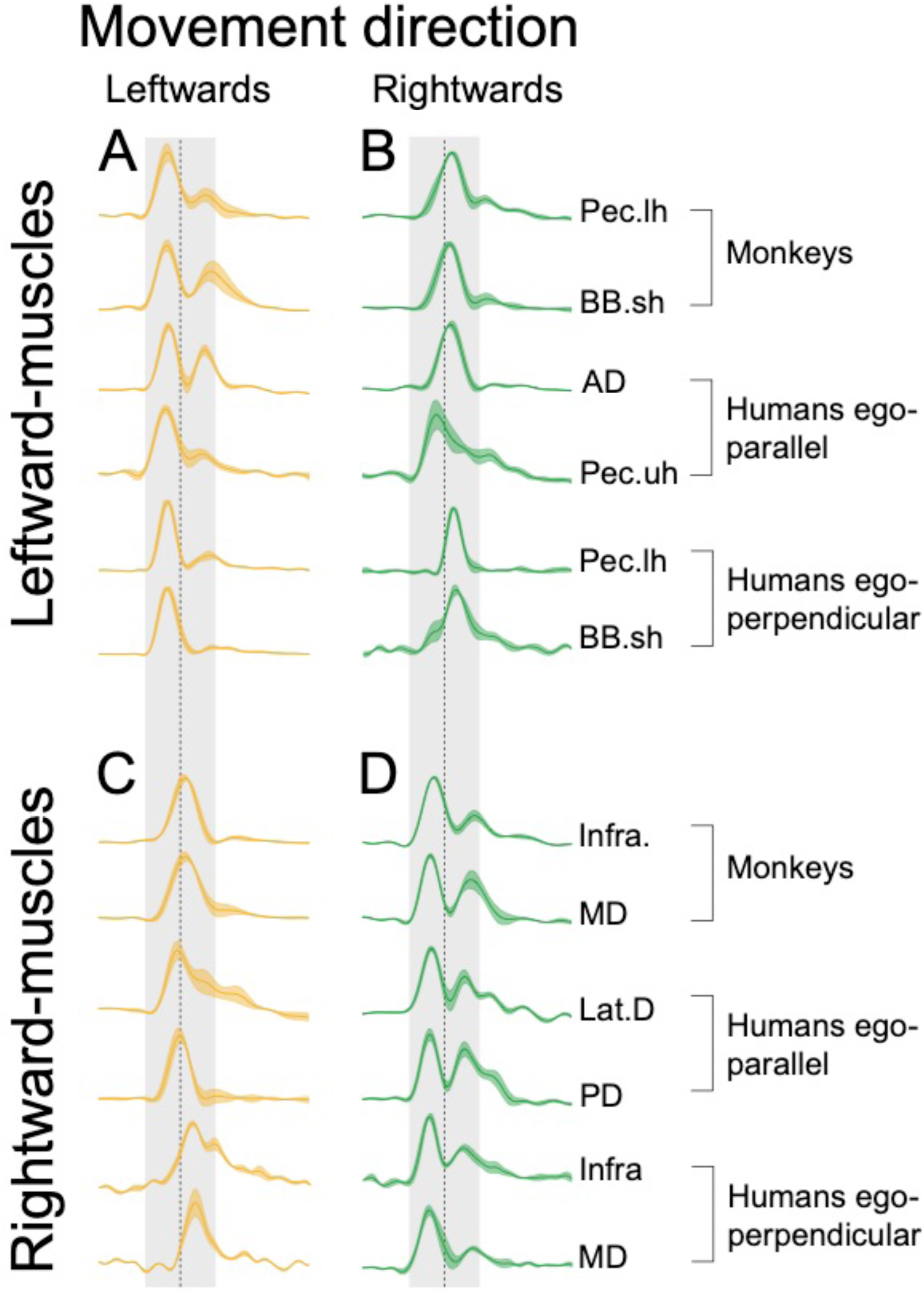
Phasic EMG patterns of earth-horizontal movements. Same layout as Figure 3. Mean (± SE) phasic EMGs are presented for monkeys (n=3) and both groups of humans (n=8 in each group). Left column (panels A and C, yellow traces) presents EMGs recorded during leftward movements, and right column (panels B and D, green traces) presents EMGs recorded during rightward movements. Upper row (A and B) presents EMG activations of muscles pulling leftwards. Those muscles pull perpendicularly to gravity, i.e. away from the final target during a rightwards movement and towards the final target during a leftward movement. Lower row (C and D) presents EMG activations of muscles pulling rightwards. Those muscles also pull perpendicularly to gravity; i.e. towards the final target during a rightward movement and away from the final target during a leftward movement. No negative periods were detected during earth-horizontal movements, meaning that gravity effects (perpendicular to movement direction) were correctly compensated. EMG traces were aligned on movement onset and normalised in duration and amplitude before averaging between participants. The grey area denotes movement duration (shifted 50 ms backwards to account for the electromechanical delay), and the dashed earth-vertical line denotes 50% of movement duration.

### Quantification of prevalence, amplitude, and duration of EMG negativity

The vast majority of movements performed in the earth-vertical plane exhibited this general muscular pattern. On average (± SD), across monkeys and humans, the phasic EMG of antigravity muscles exhibited negative values in 90.8 ± 2.7% and 72.1 ± 5.5% of downward and upward movements, respectively (see Figure 5A and Fig. 5 Suppl. Table 1 for all values). In contrast, in the earth-horizontal plane, rightwards and leftwards movements exhibited negative periods only in 0.014 ± 0.036% and 0.021 ± 0.048%, respectively (see Fig. 5 Suppl. Table 1 for values).

**Figure 5.**
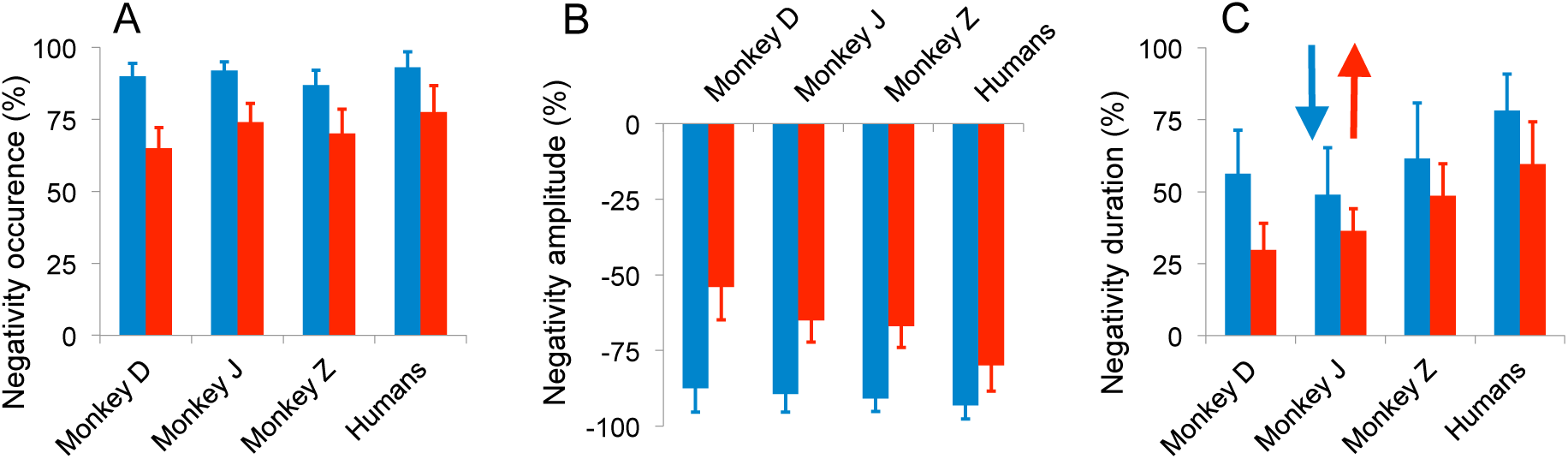
Substantial negativity of phasic EMGs. Characterisation of negative epochs recorded for antigravity muscles (the muscles pulling against gravity, presented in panels A and B of Figure 3) during earth-vertical movements. Mean values (± SD) are presented for each monkey and all humans (n=16). Coloured arrows indicate movement directions (blue for downwards; red for upwards). **A**. Mean occurrence percentage (number of positive detection/number of trial) of a negative phase for each monkey and humans. **B**. Maximal negativity amplitude was expressed as a percentage of the subtracted tonic value (see Methods). A value of −100% means the muscle was utterly relaxed. Thus gravity helped to move the arm. On the contrary, a value of 0% means gravity effects were correctly compensated for by muscle force. **C**. Mean negative phase duration expressed as a percentage of the acceleration duration for downward movement and as a percentage of the deceleration duration for downward movements. Error bars represent the standard deviation of the mean between recording sessions for each monkey and between participants for humans.

An amplitude index quantified how much gravity force assisted muscle force. This index expressed antigravity muscles activation levels relative to the theoretically required level for exact compensation of gravity effects (the tonic level; see Methods). An index value of −100% means utterly relaxed muscles, thus that the gravity force fully participated to arm movement. A value of 0% means that antigravity muscles precisely compensated for gravity effects, thus that the gravity force did not produce any movement. On average, across monkeys and humans, the amplitude index averaged (± SD) −90.7 ± 2.8% and −66.7 ± 9.6% for downward and upward movements, respectively (see Figure 5B and Fig 5 Suppl. Table 1 for all values).

We also computed the duration of negative epochs and expressed it as a percentage of acceleration duration for downward movements and as a percentage of deceleration duration for upward movements. On average (± SD), negative periods represented 60.7 ± 11.5% of downward acceleration duration and 43.1 ± 12.4% of upward deceleration duration (see Figure 5C and Fig 5 Suppl. Table 1).

In both species, the prevalence and extent of negative periods reveal that the brain did not compensate for gravity force. Instead, it lets gravity force much-assisting muscle force in producing arm movements. Next we further demonstrate this by characterizing the temporal organisation of agonist/antagonist muscular activation, which is the most basic and widely used descriptor of muscle patterns (Berardelli et al., 1996; Chiovetto et al., 2010; Cooke and Brown, 1994; Corcos et al., 1989; David et al., 2016; Forgaard et al., 2013; Huang et al., 2015; Irlbacher et al., 2006).

### Effect of EMG negativity on muscular activation timing

Figure 6 summarises the activation timings of agonist and antagonist muscles. “Agonist” and “antagonist” are generic denominations that respectively designate the muscles that pull toward or away from the final target. We found that this timing radically changes with movement direction (see Fig. 6 Suppl. Table 1 and 2 for all values and statistical comparisons). Most strikingly, it is the negativity of antagonist muscles (antigravity, open circles) that governed the acceleration of downward movements (Figure 6, blue arrow). During downward movements, both humans and monkeys activated their agonist muscles (gravity muscles, filled squares) nearly at the time of movement initiation. Given electromechanical delays (Cavanagh and Komi, 1979), this agonist activation occurs too late to explain movement initiation. The delayed activation of gravity muscles (agonist) presumably complements the negativity of antigravity muscles (antagonist) to reach the appropriate movement speed (Agostino et al., 1992). The organisation of downward movements is in sharp contrast to earth-horizontal movements, where agonist activation generated acceleration force roughly 100ms before movement onset (Figure 6, yellow and green arrows) (Berardelli et al., 1996; Cooke and Brown, 1994; Corcos et al., 1989).

**Fig 6.**
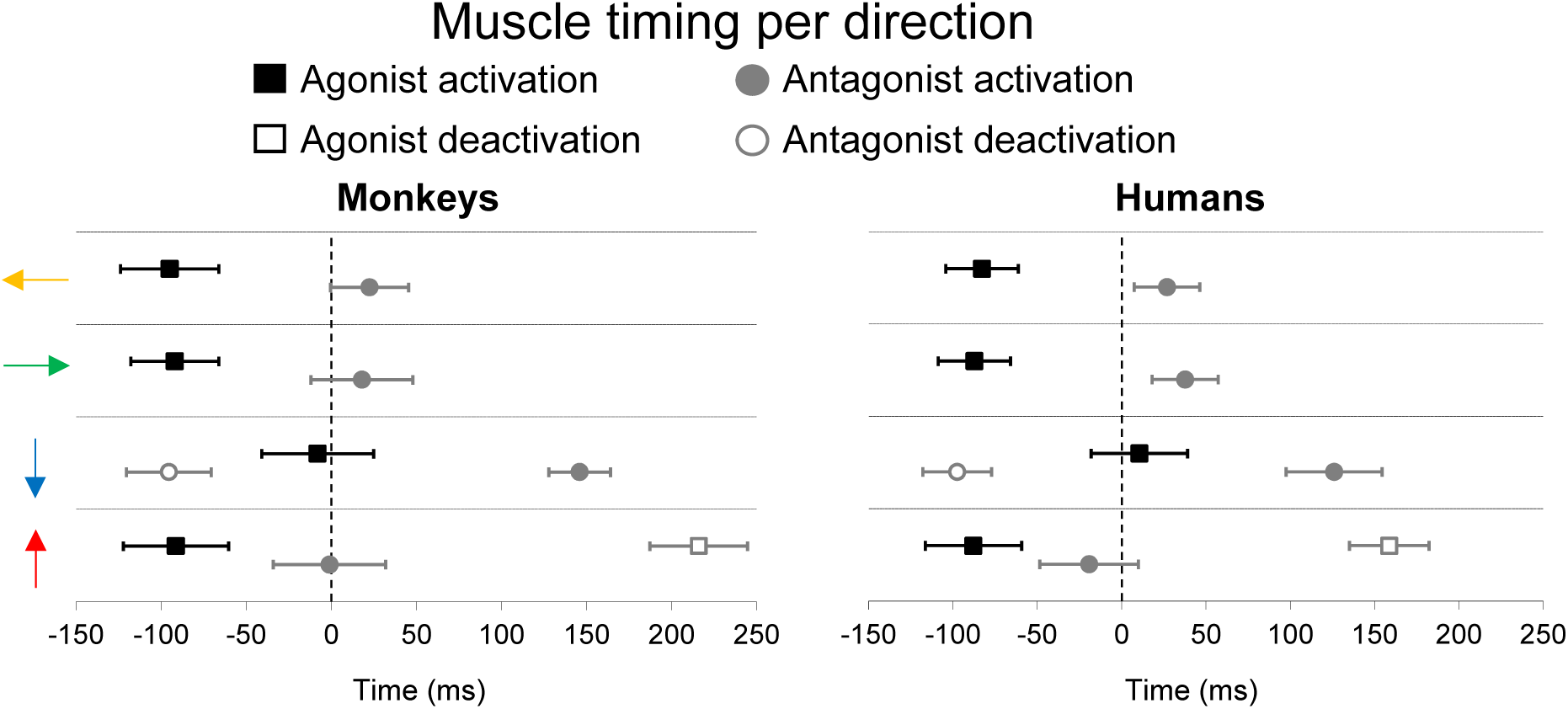
Gravity shapes the temporal organisation of muscular activation patterns. Mean (± SD) onset timings (see methods) of agonist and antagonist muscles activations for monkeys (left panel, n=3) and humans (right panel, n=16). “Agonist” and “antagonist” are generic denominations that respectively designate the muscles that pull toward or away from the final target. Muscular activity recorded in each movement direction are presented in separate rows (coloured arrows indicate movement direction). EMG traces were aligned on movement onset.

Upward movements (red arrow) exhibited an organisation that was complementary to downward movements. First, the activation of agonist (antigravity) muscles occurred at a similar timing as earth-horizontal movements (filled squares). Then, the agonist (antigravity) muscle became negative (open square). This late negativity lets gravity assist muscle force in decelerating the arm upwards. Again, this is in sharp contrast to earth-horizontal movements, where the single activation of antagonist muscles produces the deceleration force.

Thus, in summary, the temporal organisation of muscle patterns supports the *Effort*-*optimization hypothesis* according to which the brain takes advantage of gravity effects to discount muscle force that pulls downwards. Next, we directly test this hypothesis by within-muscle comparison of EMG activity between earth-horizontal and earth-vertical movements.

### Within-muscle comparison between earth-horizontal and earth-vertical movements

Humans performed arm movements that were either parallel or perpendicular to the head/feet body axis, from two body orientations (Figure 7A, B). Because the same muscles were responsible for earth-vertical and earth-horizontal movements, one can directly compare the amplitude of muscular activations between movement planes. Specifically, we will compare the activation of gravity muscles in the vertical plane to activation of the same muscles in reciprocal earth-horizontal movements (here reciprocal refers to movements that have the same direction in the ego-centred frame of reference).

**Figure 7.**
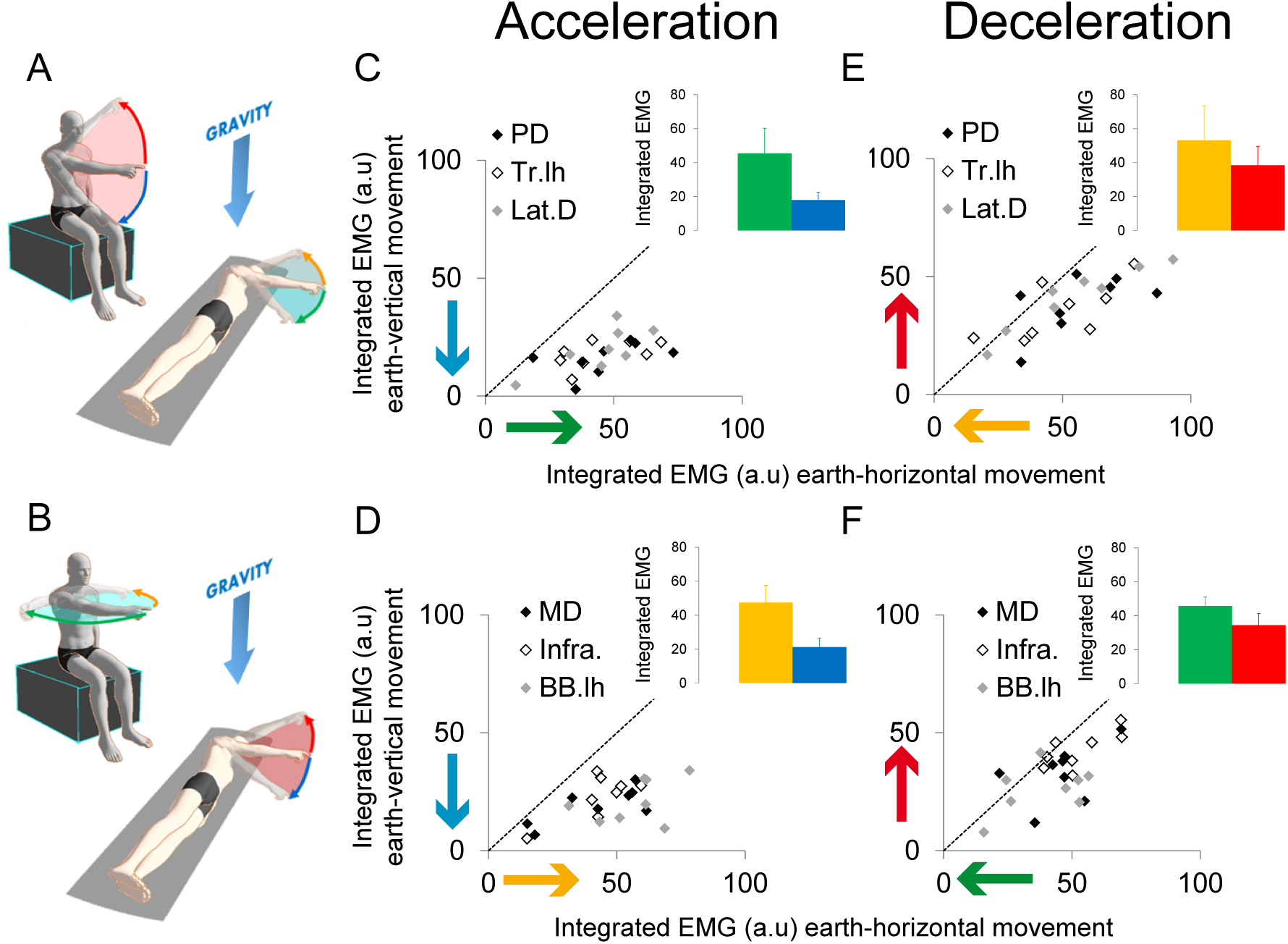
Saving effects on muscles pulling downwards. Activations of the agonist and antagonist muscles for both groups of humans (3 muscles and 8 participants per group) during earth-vertical and earth-horizontal movements. **A**. Experimental setup and movement directions in the ego-parallel group. **B**. Experimental setup and movement directions in the ego-perpendicular group. In both groups, the 90° body tilt in roll allows comparing activations of the same muscles during reciprocal movements performed in the earth-vertical and the earth-horizontal planes. Coloured arrows indicate movement and gravity directions. **C.** Ego-parallel group. For each participant (n=8) and three agonist muscles (PD, Posterior Deltoid; Tr.lh, Triceps Long Head; Lat.D, Latissimus Dorsi), the mean normalised muscular activations (± SD) integrated over the acceleration phase towards the feet are compared between body orientations. Downwards earth-vertical movements (vertical axis) are compared to rightwards earth-horizontal movements (horizontal axis). The bar-graph insert presents mean activations of all muscles and participants for the two body orientations. **D**. Ego-perpendicular group. For each participant (n=8) and three agonist muscles (MD, Middle Deltoid; Infra., Infraspinatus; BB.lh, long head of Biceps Brachii), the mean normalised muscular activations (± SD) integrated over the acceleration phase towards the left side of the body are compared between body orientations. Downwards earth-vertical movements (vertical axis) are compared to leftwards earth-horizontal movements (horizontal axis). The bar-graph insert presents mean activations of all muscles and participants for the two body orientations. **D**. Ego-parallel group. Mean normalised antagonist muscle activations (± SD) integrated over the deceleration phase towards the head are compared between body orientations. Upwards earth-vertical movements (vertical axis) are compared to leftwards earth-horizontal movements (horizontal axis). The bar-graph insert presents mean activations of all muscles (n=3) and participants (n=8) for the two body orientations. **E**. Ego-perpendicular group. Mean normalised antagonist muscle activations (± SD) integrated over the deceleration phase towards the right side of the body are compared between body orientations. Upwards earth-vertical movements (vertical axis) are compared to rightwards earth-horizontal movements (horizontal axis). The bar-graph insert presents mean activations of all muscles (n=3) and participants (n=8) for the two body orientations. The use of gravity force for downwards acceleration and upwards deceleration (as revealed by negatives phases on antigravity muscles in previous figures) results in reduced muscle activations in the earth-vertical compared to the earth-horizontal plane.

For example, in the ego-parallel group (Figure 7A), one can compare the activation of agonist muscles between acceleration phases towards the feet in the two planes (blue vs green targets). If gravity effectively assists muscle force, one would expect reduced activation of gravity muscles in the earth-vertical plane compared to reciprocal movement in the earth-horizontal plane. Following this logic, Figure 7C-F compares muscular activation between reciprocal movements performed in the two planes, for all movement phases that exhibited negativity of antigravity muscles. Data falling below the identity line indicate that earth-vertical movements necessitate weaker muscular activations than earth-horizontal ones and vice versa.

Figure 7C-D compares downward movement acceleration to reciprocal movement acceleration in the earth-horizontal plane, for the ego-parallel group and the ego-perpendicular group (blue arrows vs green arrows in Figure 7A-B). All individual comparisons fall below the identity line (n=48). This result demonstrates that downwards acceleration required reduced agonist muscle force than horizontal acceleration (all agonist muscles averaged per participant, Student paired two-sided; ego-parallel group, T_7_=6.4, P=0.0004; ego-perpendicular group, T7=9.6, P=0.00003, see bar-graph inserts in Figure 7C-D). Thus, gravity effectively assisted muscle force in accelerating the arm downwards.

Figure 7E-F compares the decelerations of upward movements and reciprocal movements in the earth-horizontal plane for the two groups (red arrows vs yellow arrows in Figure 7A-B). All but seven individual comparisons fall below the identity line (total n=48). The acceleration of upward movements required significantly reduced antagonist muscle force (all antagonist muscles averaged, Student paired two-sided; ego-parallel group, T_7_=3.3, P=0.01; ego-perpendicular group, T7=8.0, P=0.00009; see bar-graph inserts in Figure 7E-F). Thus, gravity effectively assisted muscle force in decelerating the arm upwards.

As predicted by the *Effort-optimization hypothesis*, these results confirm that the negativity of phasic EMGs effectively reduces the muscular effort that pulls downwards in gravity (not body)-coordinates. The neural integration of gravity yields optimal motor planning in both monkeys and humans.

## Discussion

Here, we investigated a fundamental question: does the motor system compensate for the gravity force, or does it exploit it as a ‘free’ force to discount muscle effort during movement? A combination of kinematics and EMG analyses have provided strong support for the *Effort-optimization* strategy. We analysed multiple variables, but two stand out as most relevant. (1) The relative time to peak velocity unmasked the motor plan at the kinematic level. (2) The negativity of the phasic EMG component revealed the signature of the optimal motor plan where gravity is purposefully exploited for effort-efficient motor control.

As previously demonstrated, shorter acceleration duration and higher peak acceleration for upward than for downward movements is the kinematic signature of optimal integration of gravity into the motor plan (Berret et al., 2008; Gaveau et al., 2014, 2016). If the motor system compensated for the gravity force, kinematics should be direction-independent in the earth-vertical plane, as is the case in the earth-horizontal plane (Gentili et al., 2007; Le Seac’h and McIntyre, 2007). The present results reveal that monkeys produce the same direction-dependent kinematics as humans (Figure 2), suggesting that optimal motor planning in the gravity field may reflect a universal cross-species process.

We sought the neural signature of this optimal motor plan by analysing EMG activity patterns. Our rationale was straightforward. If the brain truly exploits the gravity force to discount muscular effort, specific negative periods should appear in the phasic EMG signal. These negative periods would mean that the muscle produces less force than required to compensate the gravity force, such that the motor system exploits the gravity force to generate movement. We used single-degree-of-freedom movements to systematically vary the effect of gravity while the rest of the movement dynamics (inertial forces) remained constant. We found that antigravity muscles exhibited periods of negativity precisely during the acceleration phase of a downward movement and the deceleration phase of an upward movement, when gravity can assist the movement (Figure 3 & 4). Notably, many studies already observed negativity of the phasic component of muscular activations and forces during vertical movements (Buneo et al., 1994; d’Avella et al., 2006, 2008; Flanders and Herrmann, 1992; Flanders et al., 1994, 1996; Olesh et al., 2017; Russo et al., 2014). However, this phenomenon was primarily ignored and attributed to erratic errors in the separation of noisy signals. The present study demonstrates that negativity is not erratic but systematic.

Furthermore, we demonstrated that the negativity of the phasic component was both consistent and extensive (Figure 5), significantly affected the temporal organisation of the muscle patterns (Figure 6), and decreased the activation of gravity muscles (the muscles pulling with gravity, Figure 7). These results provide strong support for the active participation of the gravity force to movement generation. In both macaques and humans, direction-dependent kinematic and muscular patterns point towards a motor strategy that discounts muscle effort, as previously proposed by computational models (Berret et al., 2008; Crevecoeur et al., 2009; Gaveau et al., 2011, 2014, 2016).

Given the richness of humans’ and monkeys’ motor repertoires, the study of mono-articular arm movements may seem restrictive. However, it is essential to point out that direction-dependent motor patterns have been observed for movements as varied as mono-articular arm (Crevecoeur et al., 2009; Gaveau and Papaxanthis, 2011; Gaveau et al., 2014, 2016; Gentili et al., 2007; Hondzinski et al., 2016; Sciutti et al., 2012; Le Seac’h and McIntyre, 2007), multi-articular arm (Berret et al., 2008; Papaxanthis et al., 1998a, 1998b; Yamamoto and Kushiro, 2014) and whole-body movements (Manckoundia et al., 2006; Papaxanthis et al., 2003). These results, along with the present findings, provide conceptual support for a general theory that the brain builds internal representations of the environmental and musculoskeletal dynamics to optimise motor planning and control (Franklin and Wolpert, 2011; Guigon et al., 2008; Izawa et al., 2008; Shadmehr and Wise, 2005; Shadmehr et al., 2010; Todorov, 2004; Wolpert and Ghahramani, 2000).

Several studies have provided neurophysiological evidence of gravity internalisation in both macaques and humans (Angelaki et al., 1999, 2004; Indovina et al., 2005; Laurens et al., 2013b, 2013a, 2016; Miller et al., 2008). The present results strongly suggest that a utility of this neural representation of gravity is to discount muscle effort. This optimisation capability seems to be shared by humans and non-human primates as we observed similar kinematics and EMG patterns in both species. This cross-species neural strategy may underline the fundamental influence of gravity on the evolution, development, and function of motor systems. The metabolic rate was shown to influence body size, resource use, rate of senescence and survival probability (Berghänel et al., 2015; Brown et al., 2004; DeLong et al., 2010; Munch and Salinas, 2009; Strotz et al., 2018; Voorhies and Ward, 1999). Preserving muscle effort may thus represent an essential pursuit for the brain (Baraduc et al., 2013; Bramble and Lieberman, 2004; Carrier et al., 2011; Cheval et al., 2018a, 2018b; Farshchiansadegh et al., 2016; Huang et al., 2012; Inzlicht et al., 2018; Kurzban et al., 2013; Lee et al., 2016; Lieberman, 2015; Mazzoni et al., 2007; Morel et al., 2017; Pageaux, 2016; Pageaux and Gaveau, 2016; Selinger et al., 2015; Shadmehr et al., 2016; Walton et al., 2006; Wang and Dounskaia, 2012).

## Methods

Monkeys and humans performed standard experiments as previously described elsewhere (monkeys (Moran and Schwartz, 1999; Wang et al., 2010), humans (Gaveau and Papaxanthis, 2011; Gentili et al., 2007; Le Seac’h and McIntyre, 2007)).

### Monkey experiments

#### Setup

Three rhesus monkeys (Macaca Mullata; 5.5 to 7.2kg) participated in the study after the approval of all the experimental procedures by the Animal Studies Committee and IACUC. Monkeys were head-fixed and seated in a custom-made primate chair anchored to a virtual reality system. A mirror mounted in front of the monkey’s face at an angle of 45°, reflected the display of a monitor. Monkeys wore custom-made glasses (Kodak Wratten filters red #29 and green #61), such that visual stimulus rendered in three dimensions as red-green anaglyphs. Using an optoelectronic tracking system (NDI Optotrak Certus) the 3D position of the monkey’s right hand was fed back in real-time on the monitor as a cursor sphere (1cm radius). The monkey performed the task using his right arm. A custom-made brace was positioned on the monkey’s right arm to restrain the elbow and wrist joints, allowing motion of the arm around the shoulder joint only (Figure 1A).

#### Task

By operant conditioning, we trained three monkeys to perform fast point-to-point single degree of freedom reaching movements (shoulder rotations) between sets of two targets (1 cm radius), from an initial to a final target. We positioned the targets at arm’s length. For upward and downward movements (earth-vertical plane), we set-up two targets in a parasagittal plane crossing the centre of rotation of the animal’s right shoulder joint. We horizontally aligned the upper target with the animal’s shoulder (arm horizontal: 90° shoulder elevation, 0° shoulder abduction). We positioned the lower target such that the shoulder was extended by 20° below horizontal (70° shoulder elevation, 0° shoulder abduction). Accordingly, an upward movement consisted of a 20° shoulder flexion and a downward movement consisted of a 20° shoulder extension. For rightward and leftward movements, we set-up three targets in a transverse plane crossing the centre of rotation of the animal’s shoulders. The central-starting target was the same as the upper target for earth-vertical movements (90° shoulder elevation, 0° shoulder abduction). We also set-up two additional targets at an angle of 20° rightward (90° shoulder elevation, 20° shoulder abduction) and 20° leftward (90° shoulder elevation, 20° shoulder adduction), taking as reference the horizontal position of the arm on the central target. A rightward movement consisted of a 20° abduction (starting at the central and ending at the most rightward target), and a leftward movement consisted of a 20° adduction (starting at the central and ending at the most leftward target). Monkeys performed 200 to 300 trials per sessions (total number of trials recorded for each monkey: D=2854; J=2347; Z=2987).

#### Kinematics recording

We used an optoelectronic tracking system (NDI Optotrak Certus, 200Hz) to record the 3D position of infrared emitting diodes (markers) taped on the animal arm and brace. The most distal marker was used to provide the hand position feedback to the monkey.

#### Electromyographic recording

We recorded EMG activity with pairs of insulated single-stranded stainless steel wires (A-M SYSTEMS, 790700). Before the experiment, we inserted two twisted wires (3mm un-insulated at their ends) in each targeted muscle (1cm separation) using 33G hypodermic needles (TSK STERiJECT; for further details, see (Kurtzer et al., 2006; Moran and Schwartz, 1999)). We plugged the wires into a custom-made printed circuit board, itself linked to a differential EMG amplifier (GRASS TECHNOLOGIES, QP511). Then, to verify the appropriate positioning of each electrode, we induced muscle twitch using microstimulations. The Optotrak (ODAU, NDI) sampled the raw EMG activity at 4kHz, synchronously with kinematic signals. We recorded activations patterns of the following eleven muscles: Deltoids (Anterior, Middle and Posterior), Triceps (long and lateral heads), Biceps (two heads), Pectoralis Major (Upper and Lower), Latissimus Dorsi and Infraspinatus.

### Human experiments

Sixteen participants (4 females; mean age = 25.4 ± 5.5y; mean weight = 74.2 ± 9.6kg; mean height = 168 ± 32cm) voluntarily participated in the experiments. All participants were right-handed (Edinburg Handedness Inventory; Oldfield, 1971), with normal or corrected to normal vision and did not have any neurological or muscular disorders. The local ethics committee approved the experimental protocol that was carried out in agreement with legal requirements and international norms (Declaration of Helsinki, 1964).

#### Experimental protocol

Participants performed single-degree-of-freedom arm movements with their fully extended arm (shoulder rotations). We randomly separated the sixteen participants into two groups of eight each. Participants from the “ego-parallel” group performed arm movements in the parasagittal plane crossing their right shoulder (Figure 1B). Participants from the “ego-perpendicular” group performed arm movements in the transversal plane crossing their right shoulder (Figure 1C). For each participant, we positioned three targets in the movement plane corresponding to the respective group. For both groups, the central target implied a 90° shoulder elevation and a 0° shoulder abduction (arm perpendicular to the trunk; dark arm in Figure 1B-C). From this central target, the other two targets triggered 20° shoulder rotations that were opposite in the plane of motion (light arms in Figure 1B-C).

Participants performed arm pointing movements in two conditions of body orientation (Figure 1B-C). In one condition, they sat upright with their head-feet body axis parallel to gravity. In the other condition, they were 90° rotated in roll and lied on their left side with the head-feet body axis perpendicular to gravity. Between body-orientation conditions, target position was kept constant in the participant egocentric frame of reference (targets rotated with the participant). Half of the experiment consisted of arm movements that were parallel to the gravity vector (seating upright in the ego-parallel group, and lying on the side in the ego-perpendicular group). The other half consisted of arm movements that were perpendicular to it (lying on the side in the ego-parallel group, and seating upright in the ego-perpendicular group). We counterbalanced the order of body-orientation conditions between participants.

We instructed participants to perform accurate and uncorrected arm movements at fast speed. Before the experiment, participants performed as much practice trials as they wanted in order to familiarise with the task. Then, for each body orientation, participants performed 60 trials (30 in each direction) in a randomized order (30 trials * 2 body orientations * 2 directions * 16 participants = 1920 trials). Each trial took place as follows. The participant pointed at the central target and held it. After a random delay time (1 to 2 s), he/she received instructions about which target to point. The participant performed the movement and held the final position until the experimenter stopped the recording and informed him/her to relax.

#### Kinematics recording

We used an optoelectronic tracking system (Vicon, Oxford, UK, eight cameras, 200Hz) to record the position five reflective markers taped on the participants’ arm: shoulder (acromion), elbow (lateral epicondyle), wrist (between the cubitus and radius styloid processes), hand (first metacarpophalangeal joint), and the nail of the index fingertip.

#### Electromyographic recording

We used bipolar surface electrodes (Aurion, ZeroWire EMG, IT, 2000Hz) to record EMG activity. The GIGANET unit (Vicon, Oxford, UK) allowed synchronously recording EMG and kinematic data. We placed the electrodes on the following twelve muscles: Deltoids (Anterior, Middle and Posterior), Triceps (long and lateral head), Biceps (two heads), Brachioradialis, Infraspinatus, Pectoralis Major (Upper and lower heads), and Latissimus dorsi.

### Data analysis

We processed kinematic and EMG data using custom MATLAB scripts (MathWorks). As monkey and human data were processed similarly, we describe the data analysis for both experiments in a single section.

#### Kinematics

Kinematic data processing was similar to previous studies (Gaveau and Papaxanthis, 2011; Gaveau et al., 2014). We filtered the position (low-pass, 5Hz cut-off, fifth-order, zero-phase distortion, “butter” and “filtfilt” functions) before differentiation. A 10% threshold of the peak angular velocity defined movement onset and offset. We rejected from further analyses trials where the velocity profile presented more than one local maxima (on average < 5% of trials in monkeys and < 2% of trials in humans). We then calculated the following parameters (Fig. 2 Suppl. Figure 1A): movement duration (MD); movement amplitude; peak acceleration (PA) and relative duration to peak acceleration (rDPA= duration to peak acceleration / MD); peak velocity (PV) and relative duration to peak velocity (rDPV= duration to peak velocity / MD). We also computed angular joint displacements to control that shoulder internal/external rotations, as well as wrist or elbow rotations, were negligible.

#### EMG

We first rectified and filtered EMG signals (bandpass 20-300Hz, third-order, zero-phase distortion, “butter” and “filtfilt” functions). Then we integrated this signal over 5ms bins and cut it off 500ms before movement onset and 500ms after movement offset. To compare EMGs between muscles, participants, and datasets, we normalised each trace by the maximum value observed for the corresponding muscle in the dataset. We then averaged trials across three repetitions, resulting in 10 averaged trials to be analysed.

We used a well-known subtraction procedure that was proposed to isolate the phasic and tonic components of the full EMG signal (Buneo et al., 1994; d’Avella et al., 2006, 2008; Flanders and Herrmann, 1992; Flanders et al., 1994, 1996; Olesh et al., 2017; Prange et al., 2009b, 2012; Russo et al., 2014). We computed the average values of the integrated EMG signals from 1s to 0.5s before movement onset and from 0.5s to 1s after movement offset (Fig. 2 Suppl. Figure 2). We used these average values to compute the tonic component as a linear interpolation between them. Finally, we computed the phasic component by subtracting the tonic component from the full integrated EMG signal.

We quantified the negativity of the phasic muscular activations by calculating the following parameters: i) the duration of the negative epoch, defined as the time interval where the phasic activity dropped below zero minus the 95% confidence interval (computed on the integrated EMG signals from 1s to 0.5s before movement onset) and this for longer than 40ms; ii) an index of the amplitude of the negative epoch, computed as follows:

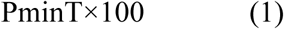

where Pmin is the phasic maximally negative value (during the negative epoch), and T is the tonic value subtracted at the time of Pmin. An amplitude index value of −100% means that muscles were completely relaxed and that the gravity force fully participated in generating the arm motion. A value of 0% means that antigravity muscles precisely compensated for gravity force; i.e., that gravity did not produce any arm motion; iii) the frequency with which a negative phase was detected amongst all trials.

We also characterized muscular activation using the following parameters: i) the onset of muscle activation, defined as the time where the phasic activity first rose above zero plus the 95% confidence interval (computed on the integrated EMG signals from 1s to 0.5s before movement onset) for longer than 40ms; v) the mean normalised integrated signal over the acceleration period (from movement onset minus 100ms to time to peak velocity minus 100ms); vi) the mean normalised integrated signal over the deceleration period (from the time to peak velocity minus 100ms to movement offset minus 100ms).

### Statistics

We checked that all variables were normally distributed (Shapiro-Wilk W test) and that their variance was equivalent (Mauchly’s test). We used repeated measure ANOVA, applied to means of separate sessions in each monkey, as well as mean values for each human subject. Post-hoc comparisons were performed using Scheffé tests. Student paired two-sided tests were used to compare muscle activation levels between body orientations in humans. In all cases, the level of significance was equal to 0.05.

## Acknowledgements

This work was supported by the Institut National de la Santé et de la Recherche Médicale (INSERM), by the Agence National de Recherche (ANR, project MOTION ANR-14-CE30-0007-01), by the Centre National d’Etudes Spatiales (CNES) and by National Institute of Health Grant R01-AT010459. We thank Daniel Moran and Thomas Pearce for their technical support.

## Figures & Legends

**Figure 2 Supplemental Figure 1.**
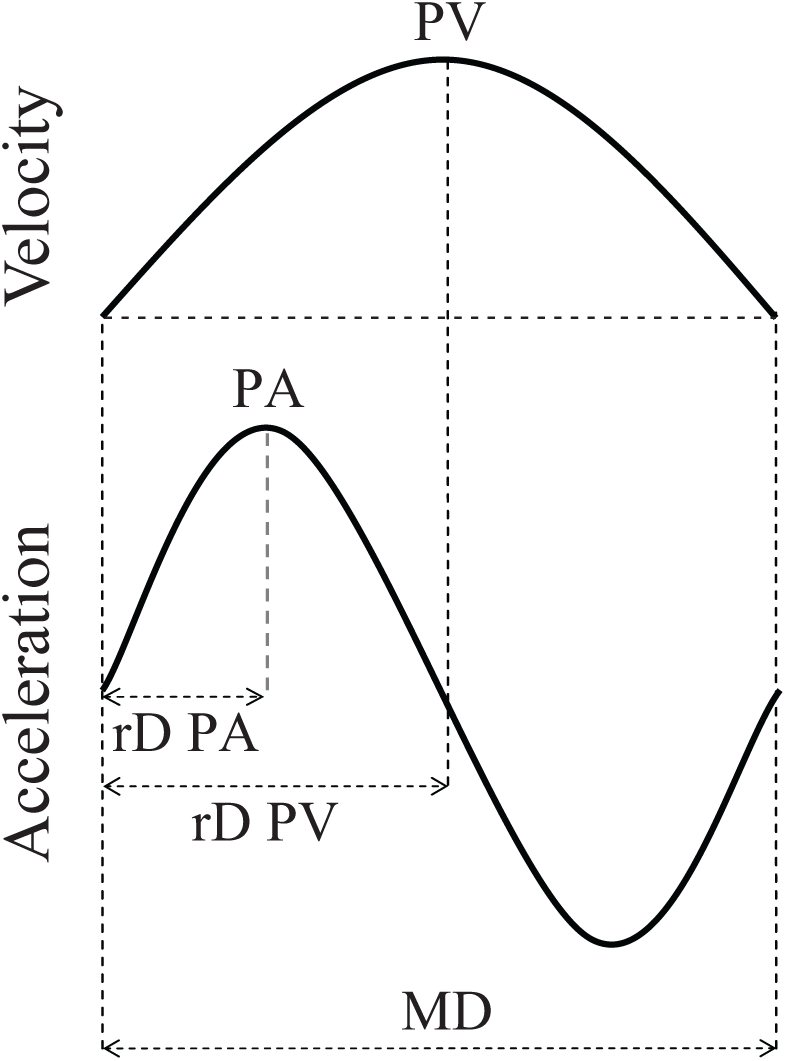
Illustration of kinematic analysis. Kinematic parameters computed on velocity and acceleration profiles (see methods). MD, movement duration; PA, peak acceleration; PV, peak velocity; rDPA, relative duration to peak acceleration; rDPV, relative duration to peak velocity.

**Figure 2 Supplemental Figure 2.**
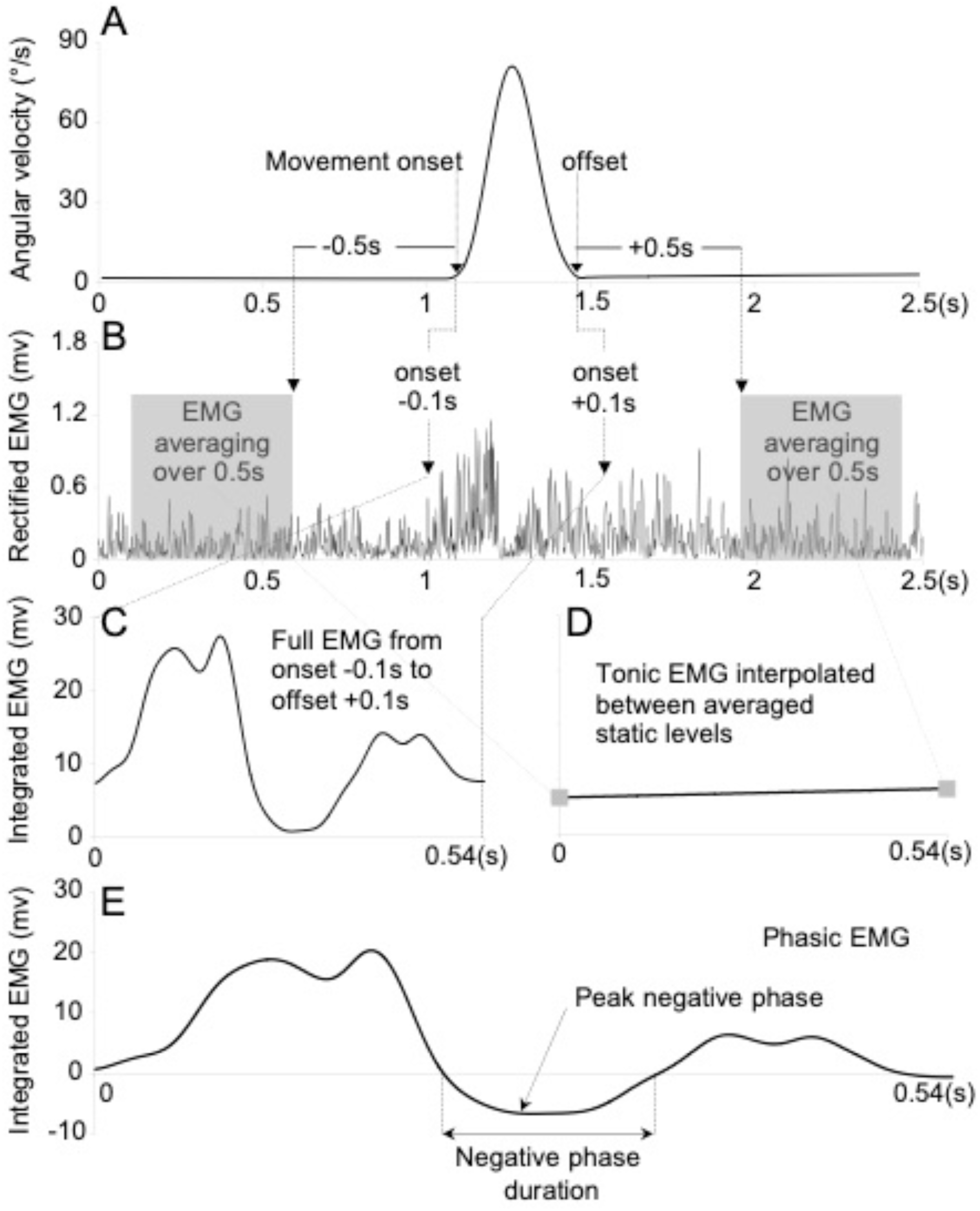
Illustration of EMG analysis. Separation procedure of the tonic and phasic component of the EMGs (see methods). **A.** Movement onset and offset were detected on the velocity profile. **B.** Timings of interest were synchronically detected on EMG data around movement onset and offset: the time intervals when the participant was waiting on the initial target (most leftward grey window) and the final target (most rightward grey window). **C.** Integrated full EMG corresponding to the movement of the arm (from 100ms before movement onset to, to 100ms after movement offset). **D.** Tonic EMG component obtained by linear interpolation between the integrated values recorded when the participant was waiting on the initial and final targets; i.e. first and last values in panel D are obtained by integrating signals from the first and second grey windows in panel B. **E.** Phasic EMG component obtained by subtracting the tonic component (D) from the full EMG (C).

**Figure 2 Supplemental Table 1.**
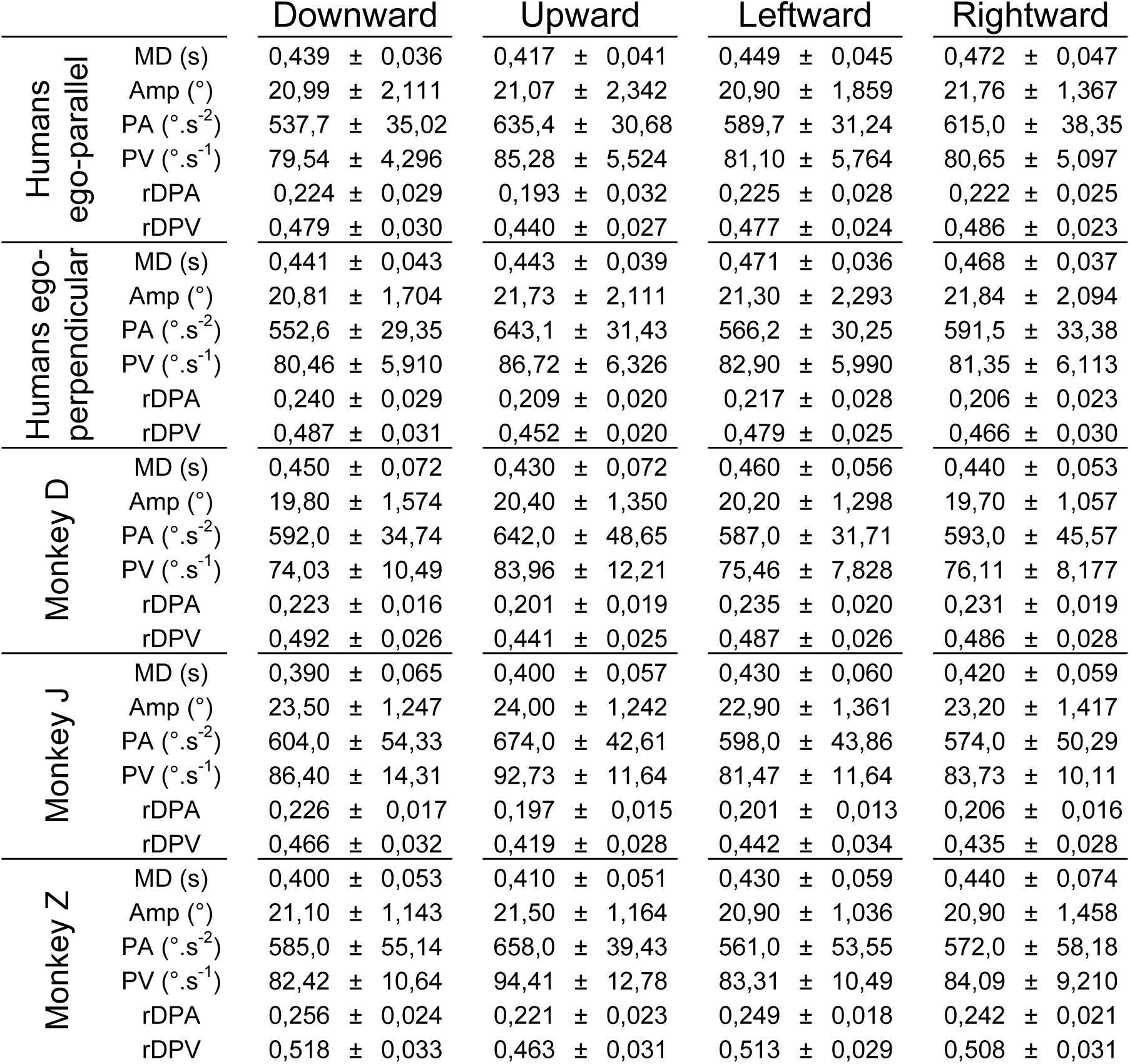
Kinematic parameters describing arm movements. Mean values (± SD) recorded for each monkey, and both groups of humans (n=8 in each group) are presented for downward, upward, leftward and rightward movements. MD, movement duration in second (s); Amp, amplitude in degree (°); PA, peak acceleration in degree per squared second (°.s^-2^); PV, peak velocity in degree per second (°.s^-1^); rDPA, relative duration to peak acceleration (duration to peak acceleration / movement duration); rDPV, relative duration to peak velocity (duration to peak velocity / movement duration).

**Figure 2 Supplemental Table 2.**
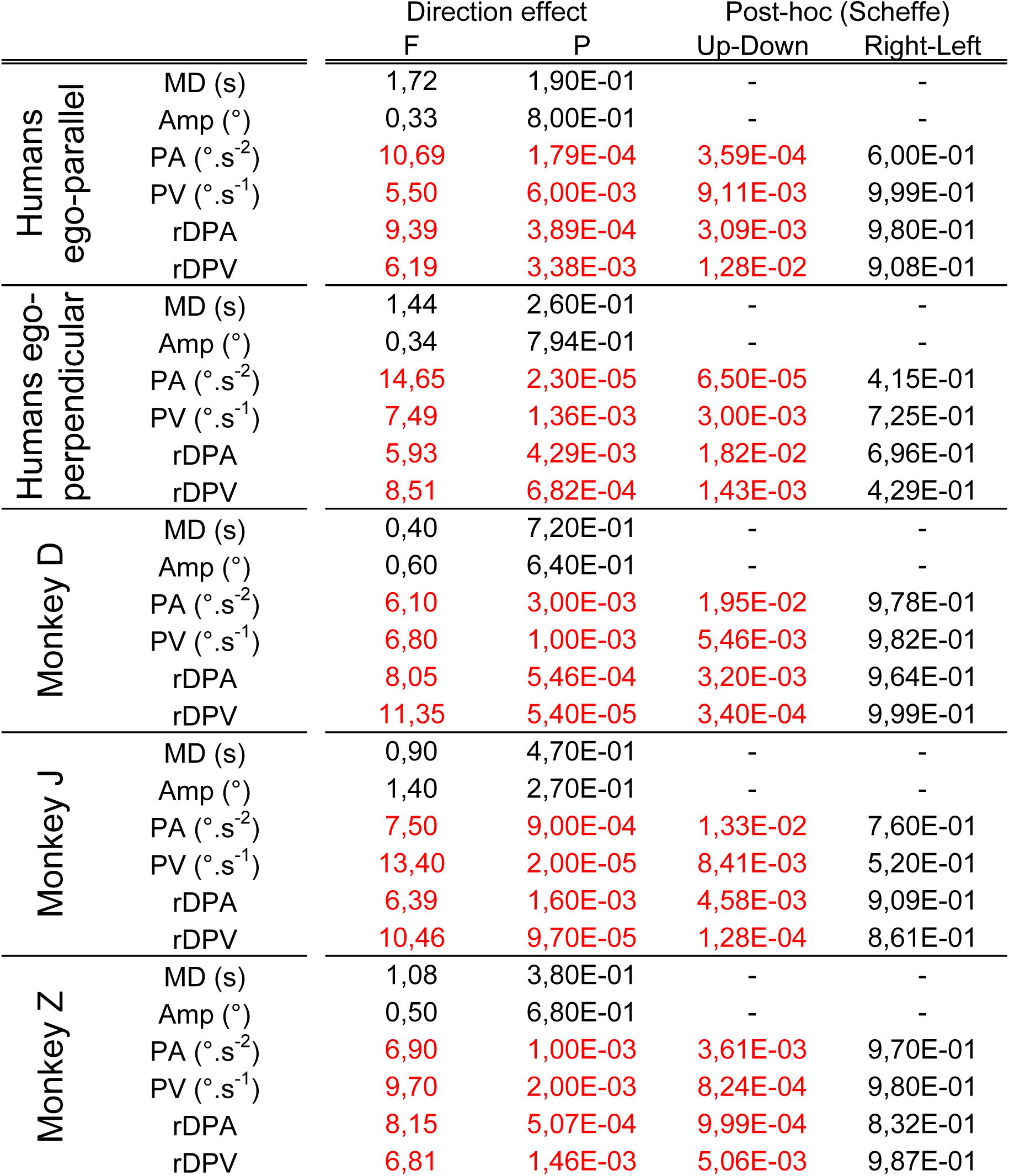
Statistical comparisons on kinematic parameters. F and P values resulting from an ANOVA test on direction effect are presented for all parameters presented in Figure 2 Suppl. Table 1. Results of post-hoc analyses (Scheffe) on opposed directions in the two planes (Up vs Down & Right vs Left) are presented for parameters on which we found a significant effect of direction (p<0.05). Like humans, monkeys exhibit directional asymmetries in the earth-vertical plane but not in the earth-horizontal plane, suggesting an optimal integration of gravity effect for the minimisation of muscular effort.

**Figure 5 Supplemental Table 1.**
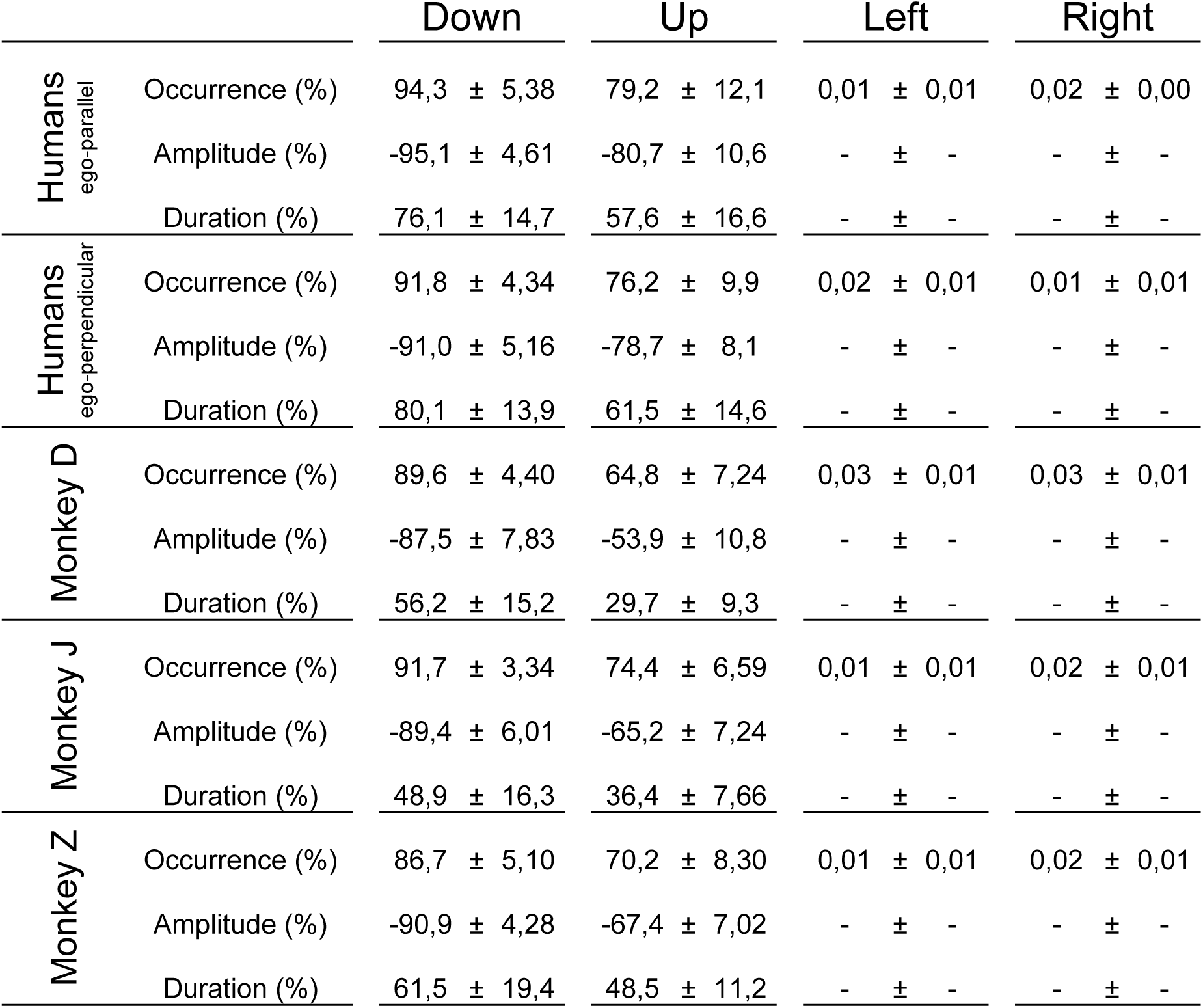
Characterisation of the negativity of the phasic component. Mean values (± SD) recorded for each monkey, and both groups of humans (n=8 in each group) are presented for downward and upward movements. Occurrence: percentage of trials where a negative phase was detected. Amplitude: percentage-ratio of negativity min value detected on the phasic component over the tonic value that was subtracted at the same time. Duration: percentage-ratio of the negative phase duration over the acceleration duration and the deceleration duration for downward and upward movements, respectively.

**Figure 6 Supplemental Table 1.**
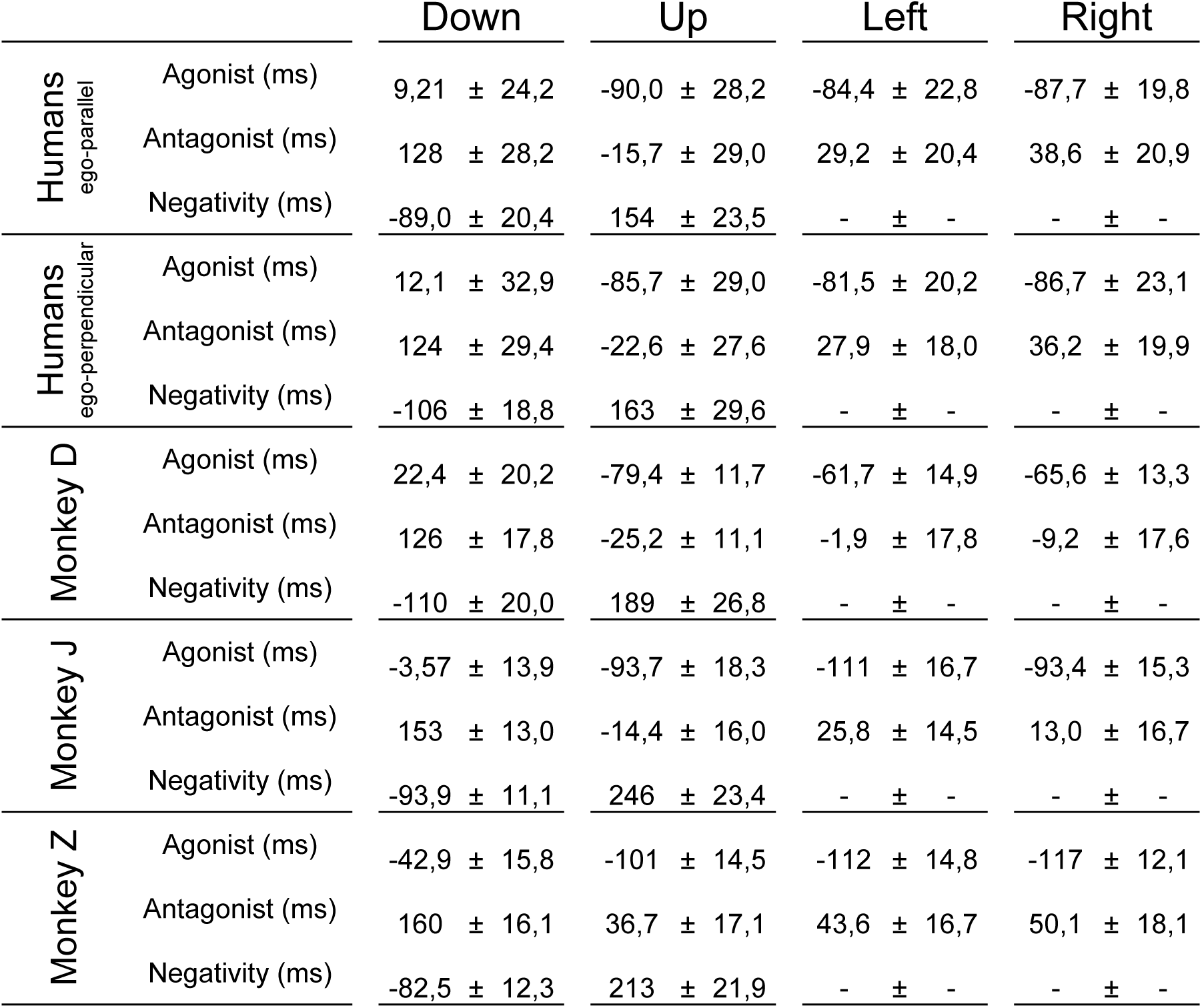
Muscle activation and deactivation. Mean values (± SD) recorded for each monkey, and both groups of humans (n=8 in each group) are presented for downward, upward, leftward and rightward movements. Timings were computed with respect to movement onset. For each direction, agonist and antagonist muscle activation timings and antigravity muscle deactivation (negativity) are provided.

**Figure 6 Supplemental Table 2.**
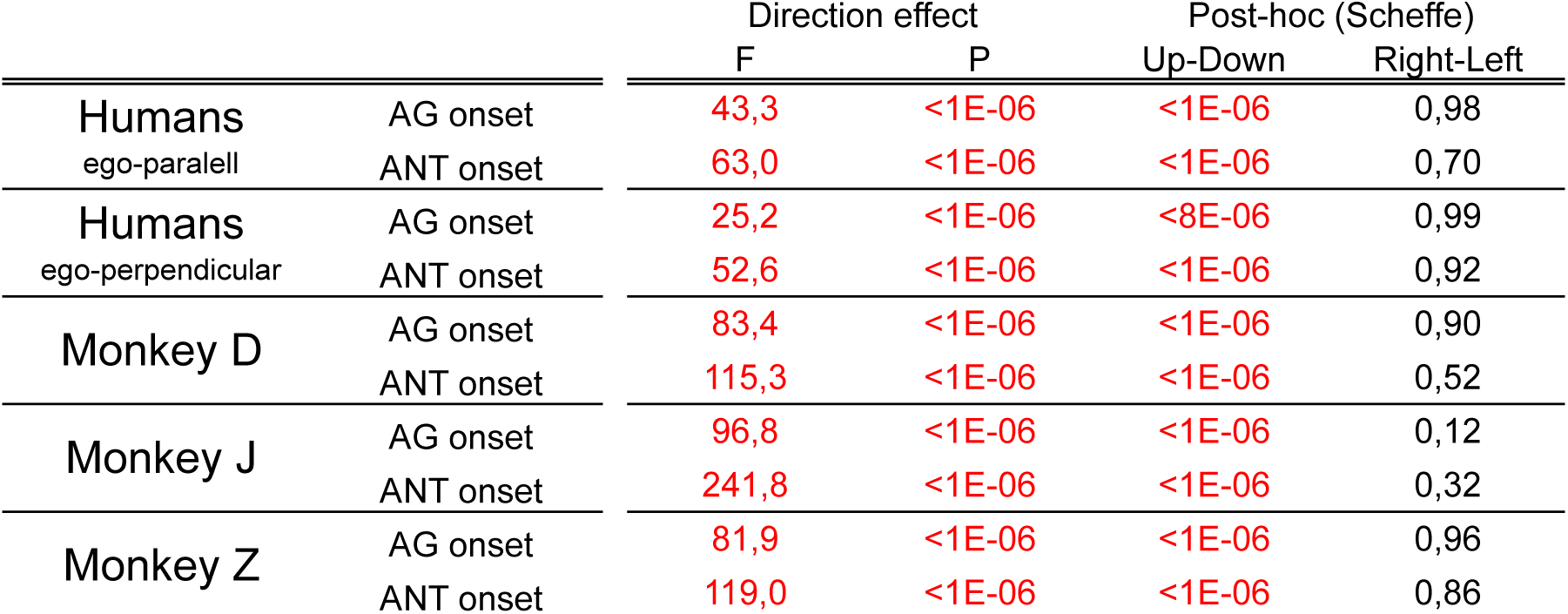
Statistical comparisons on activation timings. F and P values resulting from an ANOVA test on direction effect are presented for all parameters presented in Fig.6 Suppl. Table 1. Results of Post-hoc analysis (Scheffe) on opposed directions in the two planes (Up vs Down & Right vs Left) are also reported. Similarly to endpoint kinematics, humans (H) and monkeys (D, J & Z) exhibited directional asymmetries in the earth-vertical plane but not in the earth-horizontal plane.

